# The expression of colonic keratins is elevated in IBD, reduced in microscopic colitis, and unchanged in IBS – a retrospective study

**DOI:** 10.64898/2026.02.03.703449

**Authors:** Victor Nielsen, Lauri Polari, Emelie Lassas, Kirah Kähärä, Maria A. Ilomäki, Jannika Rovapalo, Markku Kallajoki, Markku Voutilainen, Robert J Brummer, Julia Rode, Julia König, Diana M. Toivola

## Abstract

**Background:** Keratins, a major subgroup of intermediate filament proteins, play a critical role in maintaining epithelial barrier and intracellular epithelial integrity. Studies have demonstrated possible links between inflammatory signaling and colonic keratins type II K8, and type I K18, K19 and K20, in animal models of colitis, and in patients with Inflammatory Bowel Disease (IBD). K7 is de novo expressed in patients with the IBD subtypes Ulcerative Colitis (UC) and Crohn’s Disease (CD). However, the histopathological roles of colonocyte keratins across IBD, microscopic colitis (MC), and Irritable Bowel Syndrome (IBS) remain poorly understood. Given the established utility as biomarkers in cancer diagnostics, we investigated whether keratin expression patterns could be used to distinguish inflammatory and functional colonic disorders.

**Methods:** Biobank samples from patients with IBD (n=27), MC (n=18), IBS (n=32) and healthy controls (n=31), were collected and immunohistochemically stained for K7, K8, K18, K19, and K20. Digital image analysis quantified staining intensities, which were correlated with histopathological severity scores and clinical parameters.

**Results:** Colonic keratin expression was significantly elevated in IBD, particularly in UC, while they were decreased in MC, and unaltered in IBS. Notably, K8 and K19 expression were strongly associated with areas of severe epithelial damage in IBD. Keratin expression was most pronounced in patients who had undergone colectomy due to treatment-resistant IBD.

**Discussion:** Keratin changes in IBD and MC but not in IBS highlight their importance in maintaining barrier homeostasis. Whether these changes are causes or consequences for these diseases will warrant further research.

## Introduction

Keratins (K) are intermediate filament proteins essential for maintaining the structural integrity of epithelial cells and serve as a dynamic transcellular scaffold supporting tissue architecture and function. They are classified into type I (acidic) and type II (neutral/basic) proteins, which assemble into filaments by forming heterodimers in a cell type specific manner [1].

In the human colonic epithelium, the only type II keratin is K8, and predominant type I keratins are K18 and K19. Type I K20 is also expressed, although it is absent or expressed at low levels in crypt base cells, while strongly expressed in luminal cells [2]. Keratin expression and filament assembly are dynamic and responds to cellular stress [3,4].

In the colon, keratins anchor to desmosomes, and to hemidesmosomes, thereby maintaining epithelial cohesion and resistance to mechanical stress [5,6]. This structural linkage is weakened in inflammatory bowel diseases (IBD), leading to compromised epithelial integrity and increased intestinal permeability [7]—a key feature in the pathogenesis of IBD [8]. Keratins are also linked to the nuclear lamina, establishing a structural continuum between the cytoskeleton and the nucleus. Despite these associations, the role keratins have in colonic inflammatory diseases, in which the barrier is disrupted, remains poorly understood

Chronic inflammatory conditions of the colon include inflammatory bowel disease (IBD), comprising ulcerative colitis (UC) and Crohn’s disease (CD), [9] and microscopic colitis (MC), including collagenous colitis (CC) and lymphocytic colitis (LC) [10]. These disorders share overlapping symptoms but have distinct histopathological and clinical features. Irritable bowel syndrome (IBS), although having symptomatic similarities, is classified as a functional gastrointestinal disorder with predominantly colonic symptoms, while lacking histological abnormalities in the intestinal lining [11].

Both IBS and MC appear normal in endoscopic findings. However, MC can be distinguished from IBS through microscopical analysis of biopsies. The histological hallmarks for MC are preserved crypt architecture with increased intraepithelial lymphocytes. In CC, a thickened subepithelial collagen band is observed [10,12]. These features, however, are subtle and can be challenging to assess reliably, complicating diagnosis and subtype classification [13]. Distinguishing MC from IBS has therapeutic relevance since MC is treated primarily with glucocorticoids,[14] while IBS management focuses on lifestyle interventions and symptom control [15].

In contrast to MC and IBS, IBD exhibits macroscopic mucosal changes visible during endoscopy, such as ulcers, erosions, and erythema. Microscopic examination of IBD biopsies often reveals architectural distortion of the epithelium, inflammatory cell infiltration, and, in CD, the presence of granulomas [16]. Subclassification between UC and CD is critical for treatment selection: for instance, 5-aminosalicylic acid (5-ASA) is more effective in UC [17], and colectomy may be curative in UC but is generally not curative in CD [18]. Nonetheless, diagnostic distinction can be difficult due to overlapping clinical and histological features [19], and disease severity grading remains a complex task [20]. These diagnostic challenges highlight the need for improved molecular markers to aid in the differentiation of colonic disorders. Keratins are well-established epithelial biomarkers in tumor diagnostics [21]. Altered keratin expression has been observed in IBD, with K8, K18, and K19 decreased in acute inflamed colon tissue and elevated in proximal, non-inflamed colon mucosa [22]. K7—a keratin not expressed in the healthy colon—was found to be *de novo* expressed in both CD and UC with the highest expression in areas close to ulcers [23,24]. Growing evidence indicates that intestinal barrier disruption can precede the clinical onset of IBD by several years [25] and may serve as a predictive marker for disease relapse even during periods of clinical remission [26]. This highlights the value of epithelial-derived markers—such as keratins—not only for diagnosis but also for monitoring barrier integrity and disease activity.

Here, we investigate the expression patterns of colonic keratins K8, K18, K19, and K20 in IBD and MC, and of K7—alongside K8, K18, K19, and K20—in IBS. The goal was to characterize keratin expression across a spectrum of bowel disorders in relation to tissue pathological features

## Materials and methods

### Patient material

Two cohorts of formalin-fixed, paraffin-embedded (FFPE) colon tissue samples were analyzed. Cohort I included UC (n=15), CD (n=12), CC (n=10), LC (n=8), and control samples (n=12) from Auria Biobank (Turku, Finland). The research project was authorized by the Auria Biobank’s Scientific Steering Committee (project AB17-6901) and Hospital District of Southwestern Finland (decisions T05/032/19). The same cohort was previously used to demonstrate K7 expression in IBD.^23^

Cohort II consisted of IBS samples (n=32) diagnosed by Rome IV criteria and controls (n=19) from the Örebro Biobank (Sweden; permission BA24/19). IBS subtypes included IBS-D (n=17), IBS-C (n=11), IBS-M (n=2), and IBS-U (n=2). Control tissues in both cohorts represented healthy colon mucosa, characterized by intact crypt architecture and minimal inflammatory cell infiltration. Cohort II samples were obtained from the Örebro Biobank (Örebro, Sweden) under permission number BA24/19.

In Cohort I, UC and CD samples were colectomy specimens, whereas MC and controls were colonoscopic biopsies. All Cohort II samples were mucosal biopsies from the sigmoid colon collected at a standardized location (20–25 cm from the anal verge at the crossing with the arteria iliaca communis).

### Immunohistochemistry

Immunohistochemical (IHC) staining was performed using the Ventana BenchMark Ultra automated stainer (Roche Diagnostics, Rotkreuz, Switzerland). Sections (5 μm) were stained with following primary antibodies: anti-K7 clone SP52 (Roche Diagnostics), anti-K8 clone CAM5.2 (BD Biosciences, Franklin Lakes, NJ, USA), anti-K18 clone DC10 (Abcam, Cambridge, UK), anti-K19 clone A53-B/A2.26 (Cell Marque, Rocklin, CA, USA) and anti-K20 clone SP33 (Roche Diagnostics)followed by diaminobenzidine (DAB) detection and hematoxylin counterstaining.

### Fluorescent staining and imaging

HCT-116 colon cancer cells were cultured in DMEM, supplemented with L-glutamine, penicillin – streptomycin and 10 % FBS (all reagents by Thermo Fisher, Waltham, MA, US) Prior to staining, cells were washed with PBS, detached with Accutase (Thermo Fisher), fixed with 4 % paraformaldehyde (Thermo Fisher) in PBS (pH 7.4) and permeabilized using 0.1 % Triton-X (Sigma Aldrich, St. Louis, MO, US) in PBS. Fixed cells were stained using conjugated antibody mixture (EPR17078-AF647 for K7 (abcam, Cambridge, UK); EP1628Y-AF405 for K8 (abcam), C-04-AF488 for K18 (Invitrogen, Regensburg, DE) and EP1580Y-AF647 for K19 (abcam) in PBS. Imaging was performed with Leica Stellaris 8 confocal microscope (Leica, Wetzlar, DE) and images were processed using Leica Application Suite X (Leica). The number of K7 positive cells was manually counted.

### Digital image analysis

Digital images were acquired using a Panoramic 1000 scanner (3D HISTECH, Budapest, Hungary). Epithelial cells detection and cellular DAB mean intensity for K8, K19, K19, and K20 were performed using QuPath bioimage analysis software (version 0.5.1). For K7, epithelial cells were classified as positive or negative based on a threshold corresponding to minimally detectable cytoplasmic DAB staining.

The epithelial cell layer regions of interest (eROIs) were manually annotated, excluding non-epithelial cells, with priority given to areas containing full-length, top-to-bottom crypts. For Cohort I, at least two distinct eROIs per sample were selected matching the eROIs in Polari et al [23] in parallel sections, each containing a minimum of 1,000 detectable epithelial cells. For Cohort II, one eROI per sample was selected, each with at least 500 detectable epithelial cells.

### Evaluation of inflammatory characteristics

Histopathological grading was performed on Cohort I samples by a professional pathologist as described in detail in Polari et al 2022 [23]. From both Cohort I and Cohort II, clinical information on fecal calprotectin (F-Calpro), disease duration, patient age, and BMI was collected. Clinical characteristics of patients from the cohorts are listed in Table 1. For Cohort I, reason for colectomy surgery in UC patients was also obtained.

**Table 1.**
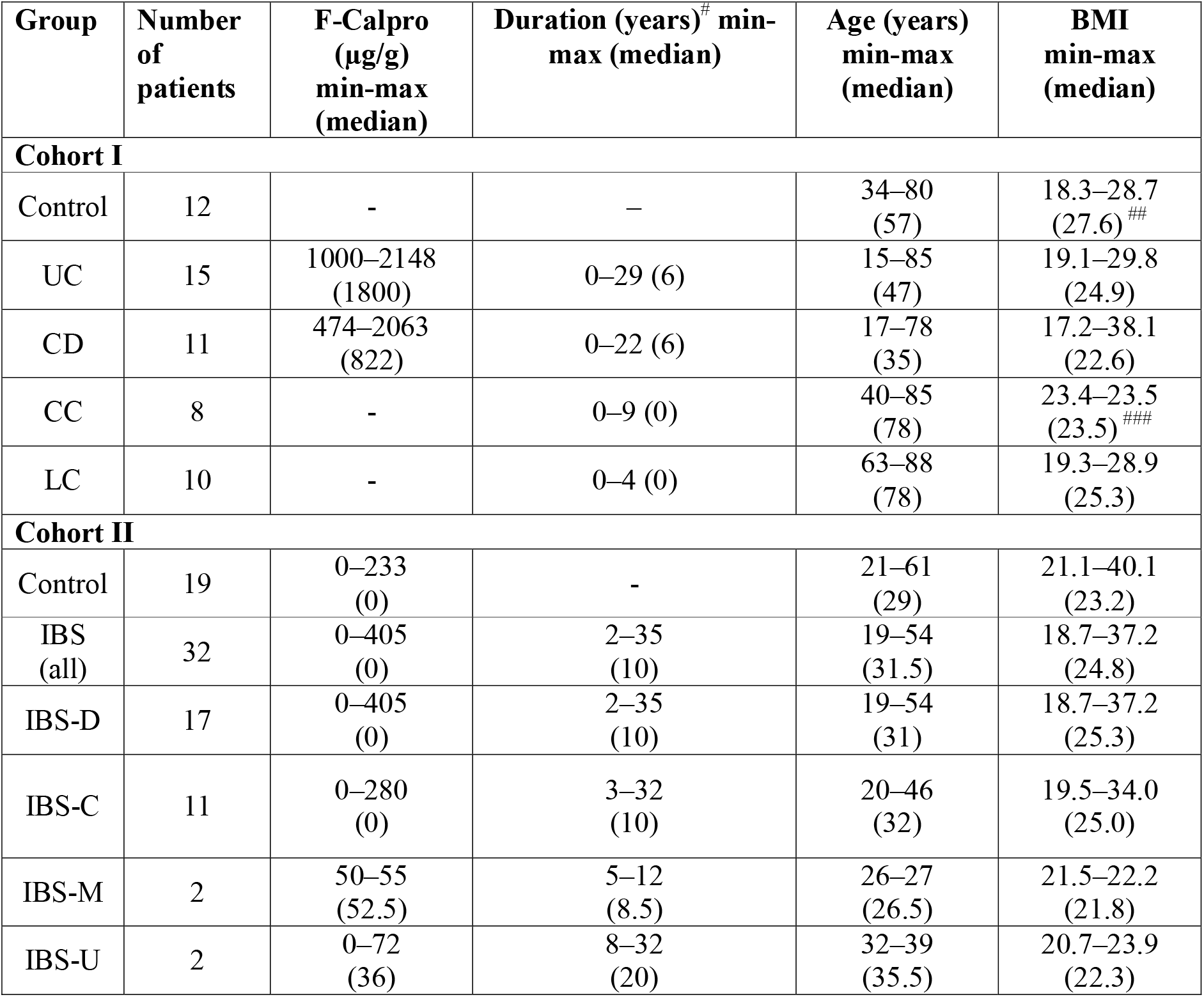
Clinical characteristics of patients. Data from Cohort I have been previously published as part of *Polari et al*., 2022 [23]. # For IBD, “Duration” refers to years since diagnosis. For IBS, “Duration” refers to self-reported years since symptom onset; ## Only 4 values available; # # # Only 2 values available. **Abbreviations:** UC, ulcerative colitis; CD, Crohn’s disease; CC, collagenous colitis; LC, lymphocytic colitis; IBS, irritable bowel syndrome.

### Statistical analysis

Statistical analyses were performed within each cohort. Differences among more than two groups were assessed using one-way ANOVA followed by Tukey’s multiple comparisons test. Comparisons between two groups were performed using the nonparametric Mann–Whitney test. Correlations between two variables were based on linear regression analysis. All statistical analyses were conducted using GraphPad Prism software (version 10; GraphPad Software Inc., San Diego, CA, USA). DAB staining intensity values were normalized to the mean value of the respective cohort-specific control group.

### Use of artificial intelligence

ChatGPT (OpenAI) was used exclusively for language editing. No scientific content was generated by the tool, and all analyses and interpretations were performed by the authors.

## Results

### Colonic keratins are upregulated in IBD and downregulated in microscopic colitis

To analyze the cellular keratin concentrations, the IHC staining DAB intensities were quantified using image analysis. In IBD, expression of K8, K18, and K19 were elevated in UC compared to controls, and K18 was also higher in CD (Fig 1A, B). Levels of K19 were higher in UC compared to CD, while K8, K18 and K20 were mostly similar between UC and CD (Fig 1B). In the MC patients, K8 was lower in LC compared to controls, and K19 was lower in CC. The decrease in keratin expression in both MC subtypes was most pronounced at the crypt base. In contrast, in both IBD subtypes, K8, K18, and K19 levels were markedly elevated throughout the entire crypts (Fig. 1A). When MC and IBD subtypes were combined into their respective disease groups, K8 was decreased in MC compared to controls. In contrast, K8, K18, and K19 were significantly elevated in the merged IBD group (Fig. 2).

**Figure 1.**
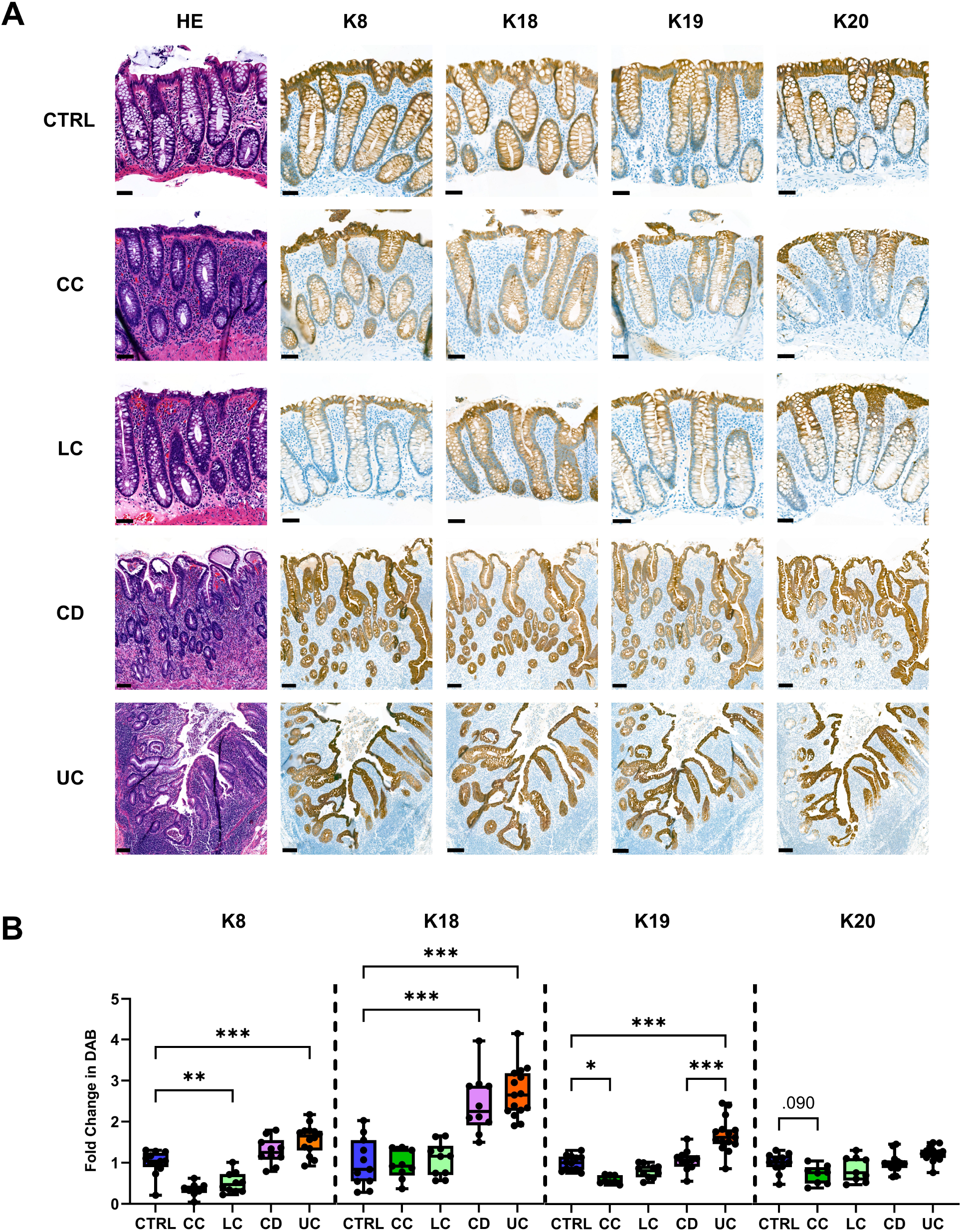
Colonic keratin staining is elevated in UC and CD and reduced in CC and LC. Formalin-fixed paraffin-embedded sections from cohort I patients were stained with HE or immunostained for keratins using IHC. **(A)** K8, K19, and K20 stainings were less intense in both MC subtypes, especially in the lower crypt regions, while the stainings of K8, K18, and K19 in both IBD subtypes were more intense throughout the crypts, compared to the normal colonic epithelium. (**B**) Cellular K8, K18, and K19 were elevated in UC as well as K18 in CD. K8 expression was lower in LC, and K19 in CC. All results in (**B**) are based on quantified cellular IHC DAB staining strength. For each sample, at least two distinct eROIs were analyzed in parallel sections, each containing a minimum of 1,000 detectable epithelial cells. Each data point represents one patient. Scale bar: control, CC, and LC = 50 μm; CD and UC = 100 μm. Statistical significance was determined after 1-way ANOVA, followed by Tukey’s multiple comparison test. Box-and-violin plots show individual data points, with the box extending from the 25th to 75th percentile and the line representing the median. Whiskers indicate the minimum and maximum values. *P < 0.05; **P < 0.001; ***P < 0.001.

**Figure 2.**
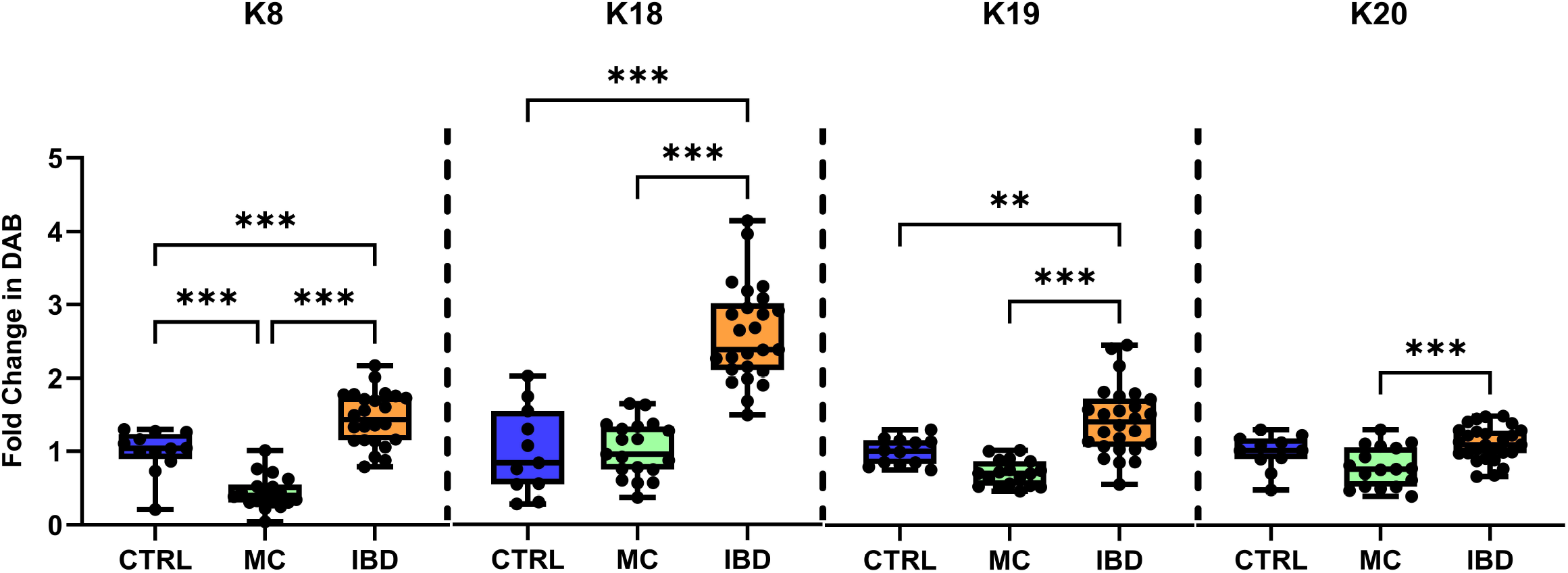
Keratin levels are increased in IBD and reduced in MC. Keratin staining data from each subtype shown in Figure 1 were merged into their respective disease groups. MC (combining LC and CC) and IBD (combining UC and CD) groups had different keratin expression patterns. K8, K18, and K19 expression were significantly higher in IBD compared to both MC and normal colon epithelium. K20 expression was also higher in IBD than in MC. Furthermore, K8 intensity was reduced in MC. Statistical significance was determined after 1-way ANOVA, followed by Tukey’s multiple comparison test. Box-and-violin plots show individual data points, with the box extending from the 25th to 75th percentile and the line representing the median. Whiskers indicate the minimum and maximum values. *P < 0.05; **P < 0.001; ***P < 0.001.

To determine which keratins are simultaneously elevated in IBD, correlation analyses were performed by regression analysis of the DAB intensity for each keratin. This analysis also included K7 DAB intensity results from a previous study, since K7 and the other keratins were done in sequential sections of the same patient samples [23]. The expression of several keratin pairs were in correlation with each other, specifically K7 and K8, K7 and K19, K8 and K19, K8 and K20, K18 and K19, and K19 and K20 (Fig. 3A, B). The results indicate that de novo expression of K7 correlates with K19, suggesting that they likely form a dimer together. This correlation was further confirmed in HCT-116 colon cancer cells in which fluorescence staining of cytoplasmic K7 fully overlapped K19 staining (Fig. 3C). K7 was also found in close proximity to K8, indicating that these two type II keratins might be present in the same filament structures (Fig. 3C).

**Figure 3.**
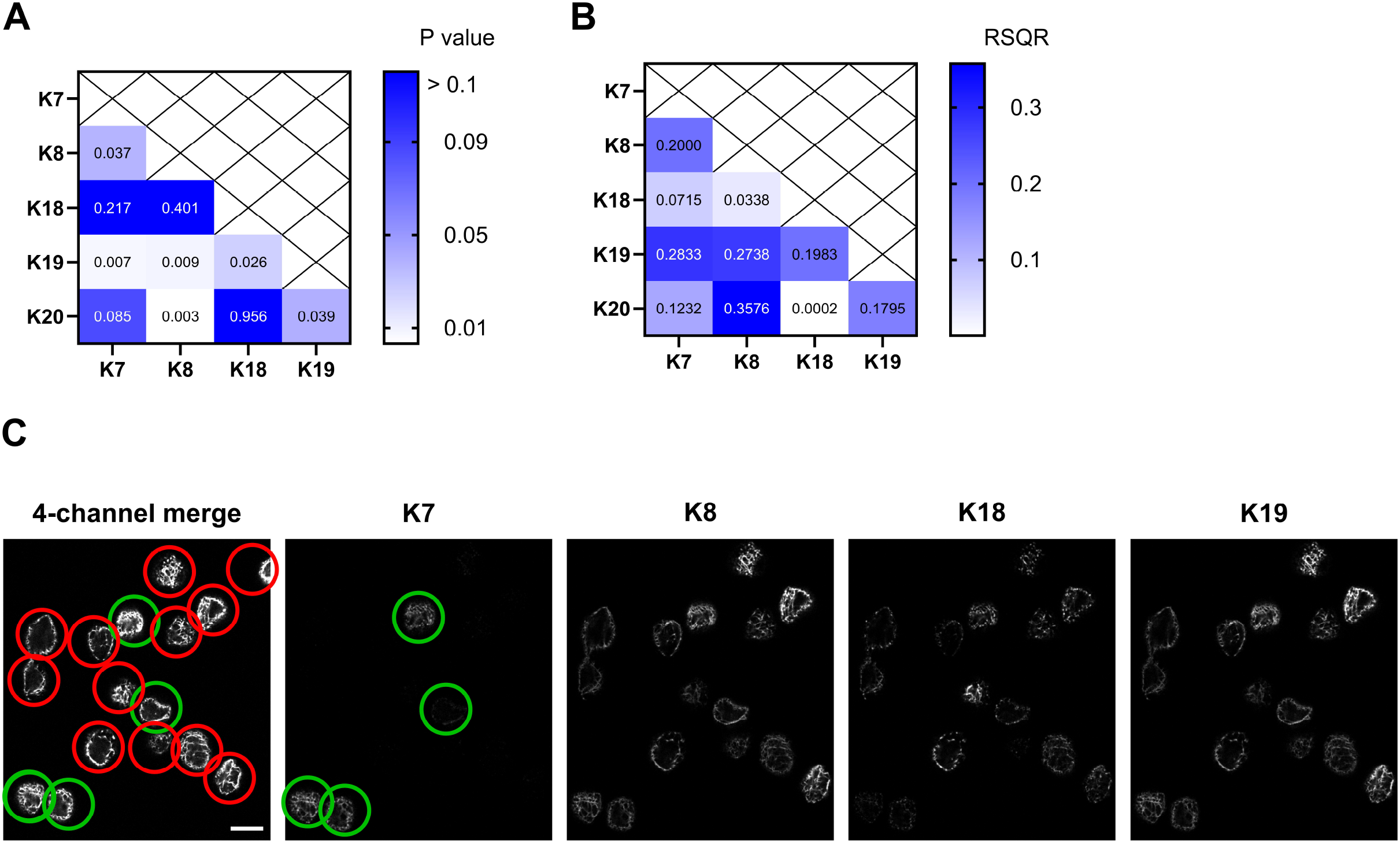
Correlation between colonic keratin expressions in IBD. Corresponding areas from adjacent sections of IBD samples in Cohort I were analyzed for (**A**) correlations between specific keratin expression using simple linear regression and (**B**) the respective RSQR values of this analysis. The analysis included K7 data from a previous study.^2^ *P < 0.05; **P < 0.001; ***P < 0.001. (**C**) HCT-116 human colorectal cancer cells were coimmunostained for keratins K7, K8, K18 and K19 using fluorophore-conjugated keratin-specific antibodies. Cytoplasmic K7 expression varied between cells and when present, it overlapped with both K19 and K8 stainings. The image on the left shows a four-channel merge of all keratin stainings while K7, K8, K18, and K19 images show respective keratin stainings. K7-positive cells are indicated by green circles (merge and K7 images) and K7-negative cells are indicated by red circles. Scale bar = 10 μm.

### Keratin expression was increased in colon of IBD patients in areas close to epithelial damage

To investigate whether spatial keratin expression correlate with IBD-related histopathological changes, two areas (aROIs) per IBD sample were evaluated by professional pathologists for key parameters, including inflammatory activity, epithelial changes, immune cell infiltration, and the presence of ulcers, erosions, or granulomas. The staining intensities of K8, K18, K19, and K20 in each the eROI of the same area (aROI) were then quantified and correlated with these histological scores.

Statistical analyses revealed significantly elevated expression of K8 and K19 in areas with severe epithelial pathologies, including deformity, atrophy, and crypt loss, compared to areas with either low or moderate changes (Fig. 4A). In addition, K8 was significantly higher in areas containing granulomas (Fig. 4D). K19 intensity was higher in areas with ulcers (Fig. 4C) and granulomas (Fig. 4D), as well as in areas scored with severe inflammatory activity compared to areas with lower inflammatory activity (Fig. 4B). None of the studied keratin intensities correlated with the high focal presence of inflammatory cells or neutrophils (Supplementary Fig. 1).

**Figure 4.**
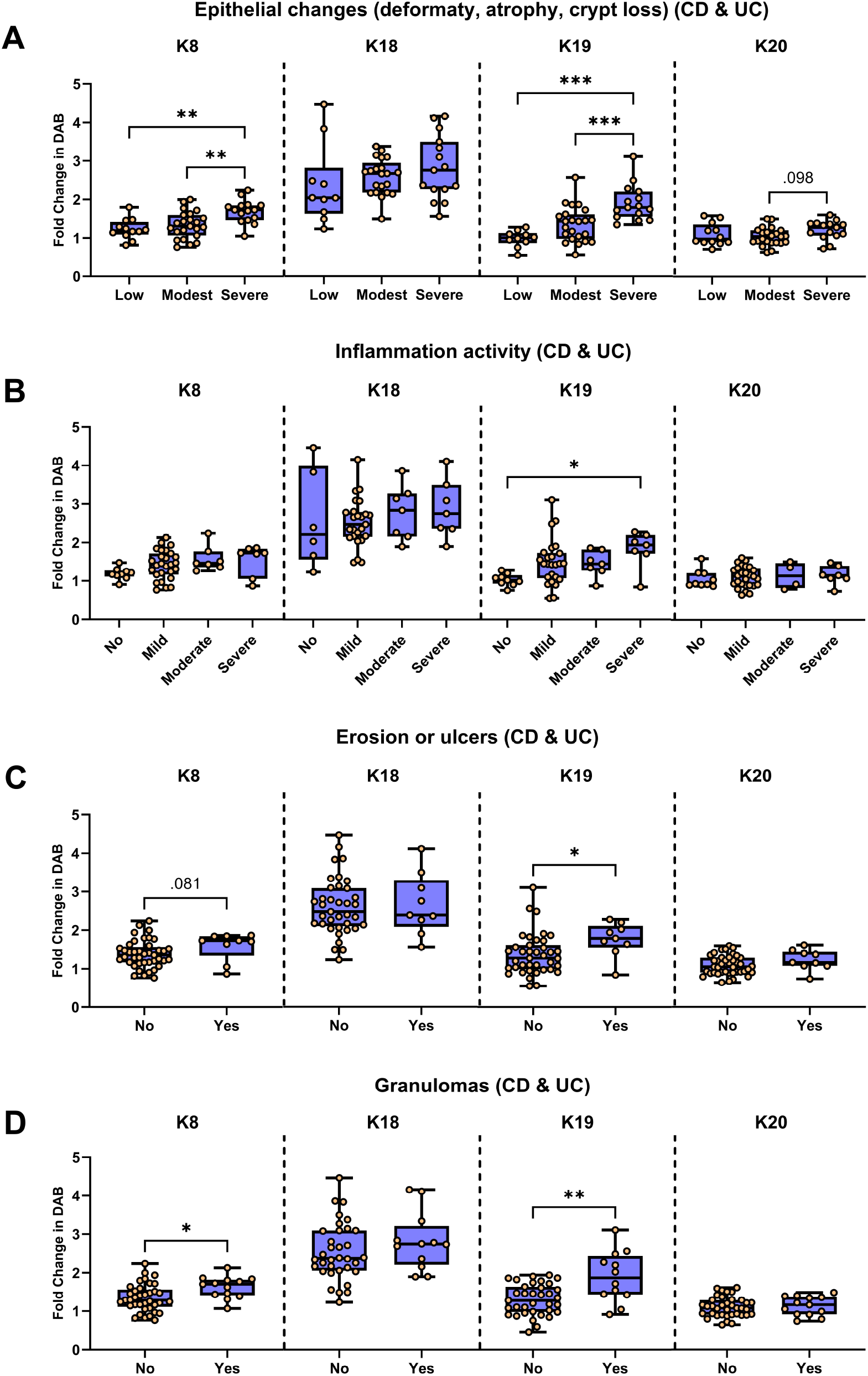
IBD-induced pathological changes are associated with elevated colonic keratin expression. Keratin expression was analyzed from two eROIs of each IBD sample in Cohort I and correlated with the grading of histopathological characteristics. (**A**) Epithelium suffering severe deformity, atrophy, and/or crypt loss had higher expression of K8 and K19 compared to areas with low or moderate changes in patients with IBD. (**B**) K19 expression was significantly elevated in close proximity to severe inflammatory activity. (**C**) Epithelial cell K19 was elevated in proximity of ulcers and erosion while K8 (p=0.081) K18 and K19 were not (**D**) Epithelial cell K8 and K19 were increased in areas with granulomas. Statistical significance between two groups was determined using the Mann-Whitney test, and comparisons among multiple groups were carried out after 1-way ANOVA, using Tukey’s multiple comparison test. Box-and-violin plots show individual data points, with each point representing one eROI. The box extends from the 25th to 75th percentile, and the line represents the median. Whiskers indicate the minimum and maximum values. *P < 0.05; **P < 0.001; ***P < 0.001.

Associations between keratin expression in IBD and clinical parameters—including the reason for colectomy in UC patients, F-Calpro, duration of IBD, patient age, and BMI—were also investigated. The results indicate reduced levels of K18 and K19 in patients who underwent colectomy due to non-drug responsiveness, compared to those who had surgery for other reasons, such as cancer, severe dysplasia, or uncontrolled inflammation, while K8 was not significantly affected (Fig. 5). No correlations were observed between the expression of any keratin and F-Calpro, disease duration, patient age, or BMI (Supplementary Table 1).

**Figure 5.**
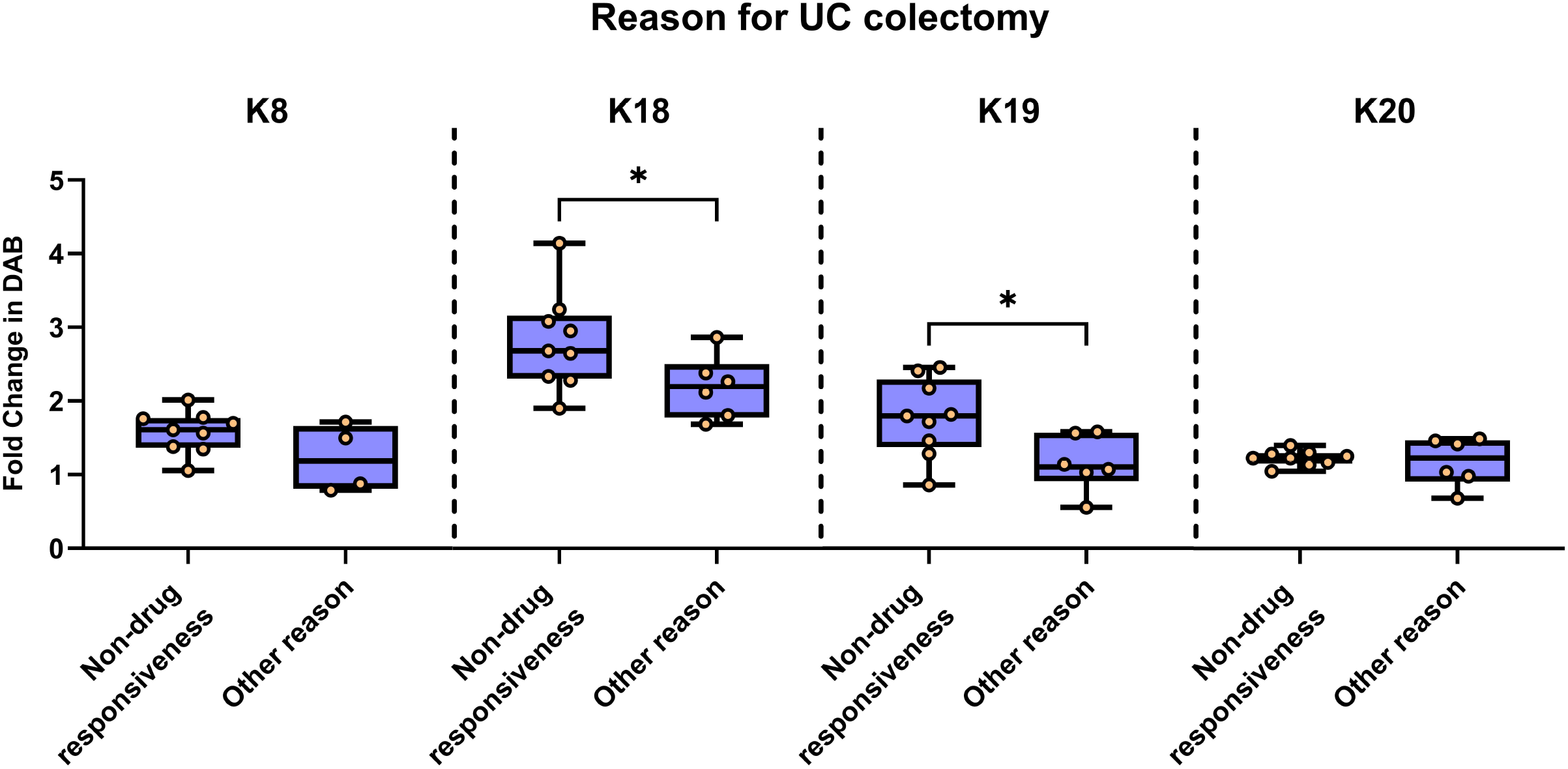
K18 and K19 levels were higher in patients that underwent colectomy due to non-drug responsiveness. Keratin expression from each IBD patient in Cohort I were analyzed against corresponding clinical data of reason the patient underwent UC colectomy (**A**) K18 and K19 levels were significantly higher in UC patients who underwent colectomy due to the disease being non-responsive to medication compared to patient that underwent surgery sue to other reasins. Statistical significance between two groups was determined using the Mann-Whitney test. Box-and-violin plots show individual data points, with each point representing one eROI. The box extends from the 25th to 75th percentile, and the line represents the median. Whiskers indicate the minimum and maximum values. *P < 0.05; **P < 0.001; ***P < 0.001.

### Keratin expression is unchanged in IBS

To investigate potential alterations in keratin expression in IBS, samples from Cohort II were analyzed for K8, K18, K19, and K20 expression, as well as K7 positivity. Results were stratified according to IBS subtype. No differences in keratin expression were observed between IBS samples and controls nor among the different IBS subtypes (Fig. 6A–D).

**Figure 6.**
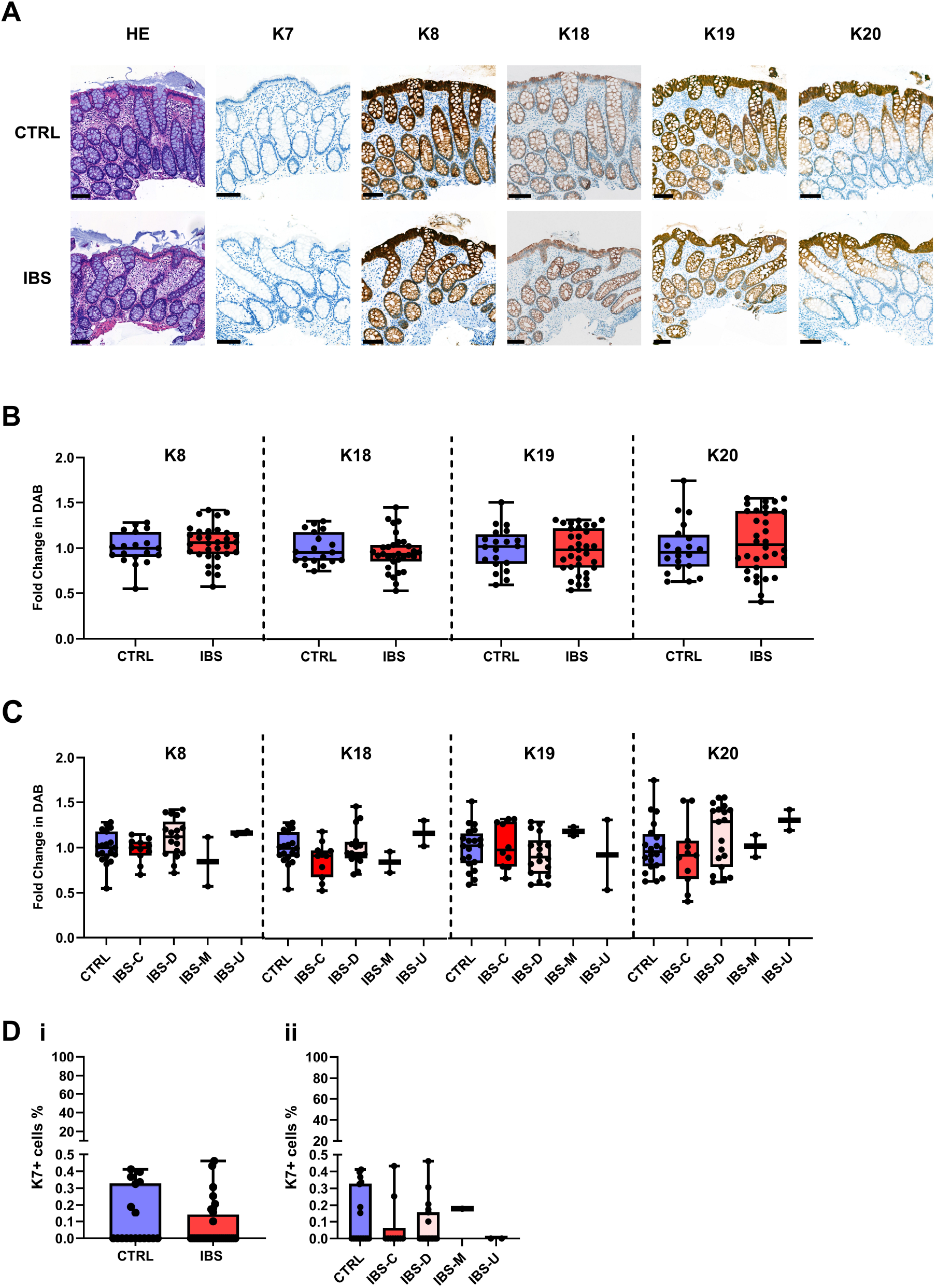
Keratin levels are unchanged in patients with IBS. Formalin-fixed paraffin-embedded sections from Cohort II patients were stained with HE or immunostained for keratins using IHC.(**A**) Cellular K8, K18, K19, and K20 levels were similar between normal and IBS colon epithelia. K7 was mostly below detection threshold in both groups. (**B**) No differences between IBS and control samples, neither when IBS subtypes were combined nor (**C**) when subgroups (C=constipation, D=diarrhea, M=mixed-type, U=unclassified) were analyzed separately. (**D**) Less than 0.5 % of the epithelial cell were K7 positive in all groups, (**i**) and there was no significant difference between combined IBS samples and normal colon epithelium, (**ii**) nor between the different IBS subtypes. Staining quantification was based on mean cellular keratin-specific DAB intensity for K8, K18, K19, and K20; and on the percentage of epithelial cells DAB stained above detection threshold for K7. One eROI per sample was analyzed, containing at least 500 detectable epithelial cells. Each data point represents one patient. Scale bar = 100 μm. Statistical significance between two groups was determined using the Mann-Whitney test, and comparisons among multiple groups were performed after 1-way ANOVA using Tukey’s multiple comparison test. *P < 0.05; **P < 0.001; ***P < 0.001.

Staining patterns in IBS tissues were identical to those in controls: K8, K18, and K19 were detected throughout the crypts, while K20 was absent or expressed at low levels in the crypt base and strongly expressed in the luminal epithelium (Fig. 6A–C). K7 expression was not detected in epithelial cells of any IBS samples (Fig. 6A & Di–ii). Rare K7-positive unidentified cells were observed within the epithelium in all samples (Fig. 6A).

A significant correlation between K8 expression and F-Calpro was observed in the IBS-C subgroup. Additionally, a near-significant association was observed between K18 expression and F-Calpro in the merged IBS cohort (p = 0.056). However, both correlations appeared to be driven by single outlier values. No other significant correlations were found between keratin expression and clinical parameters, including F-Calpro, IBS duration, age, or BMI, either when all IBS patients were analyzed combined or when stratified into IBS-C and IBS-D subgroups (Supplementary Table 2).

## Discussion

The major finding of this study is that the two main simple epithelial keratins in the colon epithelium —type II K8 and type I K19 [2]—are upregulated in IBD lesions exhibiting a higher degree of crypt distortion. This observation is consistent with previous proteomic data showing increased levels of K8, K18, and K19 in patient with chronic inflammation compared to healthy colonic tissue. Notably, the same study reported decreased levels of these keratins in cases of acute inflammation [22]. Additionally, quantitative image analysis in our study revealed reduced levels of keratins K8, K19, and K20 in MC, consistent with a proteomic study showing decreased protein expression of K8 and K20 in both MC subtypes [27].

Since keratins are well-established protectors of epithelial barrier integrity, this reduction may be linked to the cytokine-driven epithelial barrier disruption observed in MC [28]. [29]On the other hand, IBD is characterized by altered immune activation and subtle or absent symptoms prior to clinical diagnosis. The elevated keratin expression observed in IBD may correspond to the chronic nature of the disease, involving prolonged epithelial stress and remodeling during its extended preclinical phase [30].

Unlike IBD and MC, no keratin changes were found in IBS samples. Although increased gut permeability in IBS has been linked to dysregulation of tight junction proteins [31], this does not appear to involve alterations in keratin expression. This further supports the understanding of IBS— as defined by the Rome IV criteria—as one of the disorders of gut–brain interactions (DGBI) [32] with only limited immune activation [33].

Histologically, active colitis is characterized by neutrophil infiltration within the epithelium, presence of ulcers or erosions, and otherwise generally preserved epithelial architecture in the colon. In contrast, chronic colitis more commonly features lymphocytic aggregates, granuloma formation, and architectural distortion or atrophy of the epithelium [34]. These distinct histopathological features align with our correlation analyses, which revealed that keratin expression—particularly of K8 and K19—is most strongly upregulated in regions exhibiting crypt abnormalities typical of chronic inflammation. In contrast, keratin expression was not significantly increased in areas with major neutrophil infiltration in the lamina propria or epithelium. However, K18 and K19 were elevated in areas close to ulcers or erosion and modestly increased in inflamed areas.

Our findings are consistent with both in vivo and in vitro studies demonstrating that K8 and K19 play essential roles in tissue regeneration and inflammatory response. In vivo studies in mice have shown that K19-positive colonic stem cells exhibit greater resistance to inflammatory injury compared to their K19-negative counterparts [35]. Moreover, K19-positive cells actively contribute to epithelial regeneration following inflammation-induced damage, whereas K19-negative cells do not significantly participate in this reparative process [36]. Collectively, these findings suggest that overall keratin upregulation in IBD is predominantly associated with chronic epithelial remodeling. Further investigation into the spatial and temporal dynamics of keratin expression may provide additional insight into epithelial adaptation during different phases of colitis.

With respect to K8, studies in mice heterozygous for K8 deletion have shown increased susceptibility to dextran sulphate sodium (DSS)-induced colitis compared to wild-type controls. These heterozygous mice exhibited elevated histological markers of inflammation—such as erosion, crypt loss, crypt abscesses, and lymphoid aggregates [37]—closely resembling the pathological features observed in IBD. Together these findings suggest that sufficient K8 expression is required to maintain colonic epithelial integrity and to limit inflammation-associated damage. Supporting this, an in vitro study using human colorectal epithelial cells showed that K8 and K19 levels increased in response to induced inflammatory stress, whereas K18 and K20 did not [3]. This reinforces the idea that K8 and K19 are selectively upregulated during inflammation and may serve as key responders to epithelial stress.

In conclusion, the distinct expression patterns of colonic keratins observed in our study—namely the upregulation of K8, K18, and K19 in IBD, and the downregulation of K8, K19 in MC—suggest that during active, diagnosable disease, the molecular milieu in epithelial cells is considerable different. This may enable the development of novel diagnostics to distinguish between IBD, MC and IBS. In addition, results here support the biomarker potential of K7 in IBD, as K7 was not detected in the colon of IBS patients. Nevertheless, further studies analyzing their levels in tissue, stool, and serum are necessary to establish clinically relevant thresholds.

## Acknowledgements

This research was supported by the Academy of Finland (315139/332582), including the InFLAMES Flagship Programme (337531); Novo Nordisk Foundation (NNF23OC0087039); the Sigrid Juselius Foundation; ÅAU Center of Excellence of Cellular Mechanostasis; Medicinska Understödsföreningen Liv och Hälsa Foundation; Business Finland; the Tor, Joe, and Pentti Borg Memorial Fund; Satakunta Hospital District, Pori, Finland; the Swedish Cultural Foundation; and the Satakunta Regional Fund of the Finnish Cultural Foundation.

We thank Auria Biobank (Turku, Finland) for sample gathering, tissue sectioning, and H&E staining for Cohort I; the Department of Pathology in Pori for IHC and the TYKS pathology lab for IHC staining support; and Richard Forsgård for facilitating collaboration with Örebro University for Cohort II.

## Tables

**Supplementary Table 1.** Correlation of clinical parameters with keratin expression in IBD. Keratin expression from each IBD patient in Cohort I were analyzed against corresponding clinical data, F-Calpro, disease duration in years, age and BMI. Analyses were performed separately for each IBD subtype. o significant correlations were observed between the expression of K8, K18, K19, or K20 and patient age, BMI, F-Calpro, or disease duration. Correlations between two variables were assessed using simple linear regression analysis. **Abbreviations:** CD, Crohn’s disease; UC, ulcerative colitis; F-Calpro, fecal calprotectin; R2; coefficient of determination; p, probability value.

| **CD** |  | **K8** |  | **K18** |  | **K19** |  | **K20** |  |
| --- | --- | --- | --- | --- | --- | --- | --- | --- | --- |
| n=11 |  | R<sup>2</sup> | p | R<sup>2</sup> | p | R<sup>2</sup> | p | R<sup>2</sup> | p |
|  | F-Calpro# | 0.000 | 0.960 | 0.051 | 0.559 | 0.000 | 0.993 | 0.098 | 0.412 |
|  | Duration | 0.041 | 0.551 | 0.110 | 0.319 | 0.021 | 0.670 | 0.011 | 0.759 |
|  | Age | 0.004 | 0.859 | 0.027 | 0.632 | 0.014 | 0.727 | 0.003 | 0.878 |
|  | BMI | 0.023 | 0.658 | 0.046 | 0.528 | 0.206 | 0.160 | 0.001 | 0.916 |
| **UC** |  | **K8** |  | **K18** |  | **K19** |  | **K20** |  |
| n=15 |  | R<sup>2</sup> | p | R<sup>2</sup> | p | R<sup>2</sup> | p | R<sup>2</sup> | p |
|  | F-Calpro | 0.000 | 0.697 | 0.051 | 0.333 | 0.000 | 0.866 | 0.098 | 0.840 |
|  | Duration | 0.024 | 0.585 | 0.045 | 0.446 | 0.035 | 0.502 | 0.150 | 0.153 |
|  | Age | 0.028 | 0.555 | 0.065 | 0.359 | 0.048 | 0.433 | 0.000 | 0.942 |
|  | BMI | 0.086 | 0.354 | 0.020 | 0.663 | 0.015 | 0.706 | 0.150 | 0.216 |

**Supplementary Table 2.**
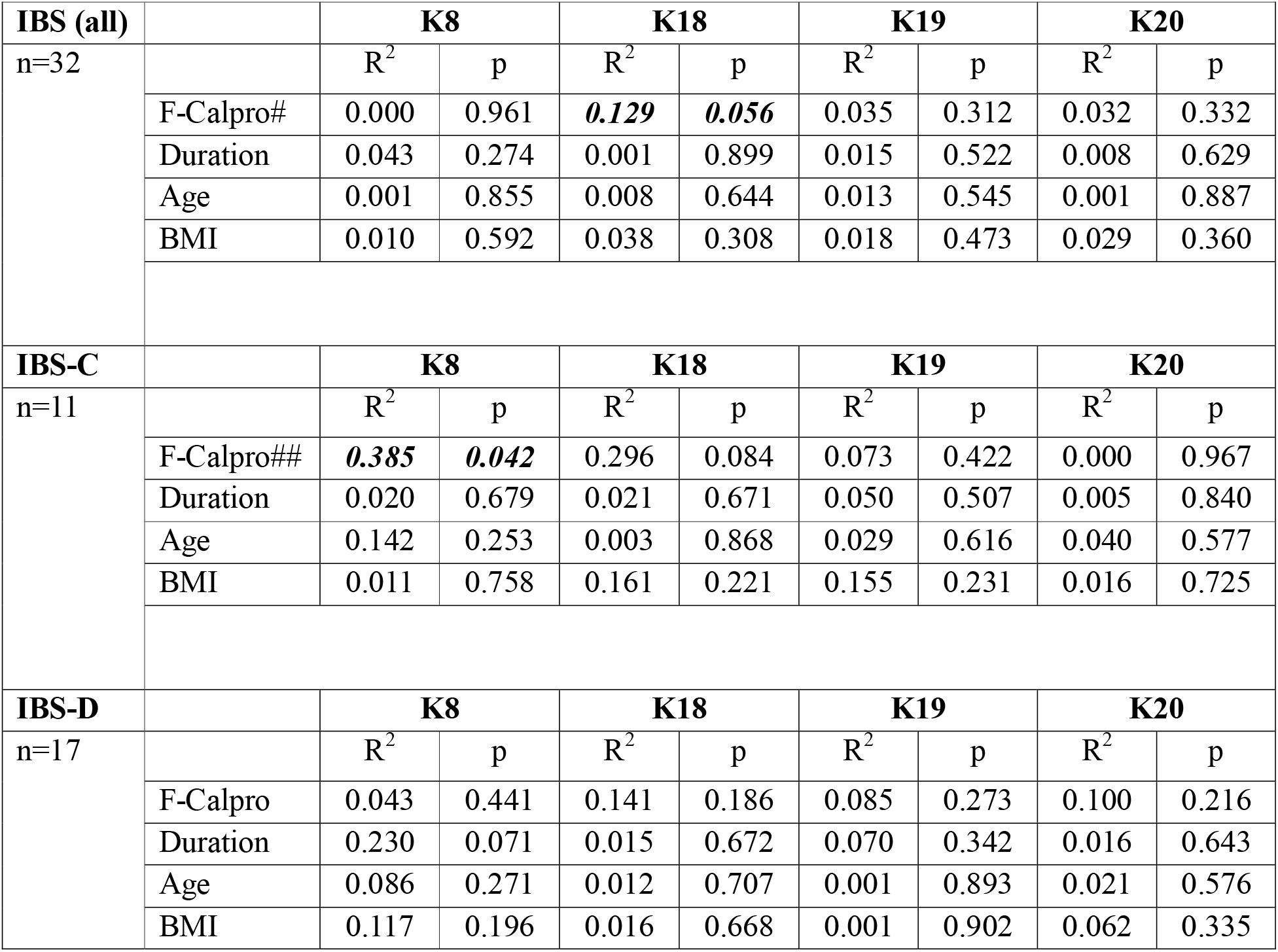
Correlation of clinical parameters with keratin expression in IBS. Keratin expression from each IBS patient in Cohort II were analyzed against corresponding clinical data, F-Calpro, disease duration in years, age and BMI. Most of the correlations were non-significant between K8, K18, K19, and K20 expression, and patient age, BMI, F-Calpro, and duration of IBS, either when all IBS cases were analyzed together or when stratified into IBS-C and IBS-D subgroups. Correlations between two variables were assessed using simple linear regression analysis. *P < 0.05; **P < 0.001; ***P < 0.001.#Positive correlation for F-Calpro dependent on a single high-value outlier (n = 11); # #Only 4 F-Calpro-positive cases, correlation driven by one high-value outlier. **Abbreviations**: IBS, irritable bowel syndrome; IBS-C, irritable bowel syndrome with constipation; IBS-D, irritable bowel syndrome with diarrhea; F-Calpro, fecal calprotectin; R2; coefficient of determination; p, probability value.

## Figure legends

**Supplementary Figure 1. No significant correlation between colonic keratin expression and
the presence of inflammatory cells**.Keratin expression was analyzed from two eROIs of each IBD sample in Cohort I and correlated with the grading of inflammatory cell presence. **(A**) Keratin expression was not affected by the number of inflammatory cells in close proximity to epithelia, (**B**) neutrophil infiltration in the lamina propria, or (**C**) neutrophil infiltration in the epithelium, compared to areas with less increased inflammatory cells or neutrophil infiltration. Statistical significance was determined after 1-way ANOVA, followed by Tukey’s multiple comparison test. Box-and-violin plots show individual data points, with the box extending from the 25th to 75th percentile and the line representing the median. Whiskers indicate the minimum and maximum values. *P < 0.05; **P < 0.001; ***P < 0.001

