## Supplementary figures and images for "The expression of colonic keratins is elevated in IBD, reduced in microscopic colitis, and unchanged in IBS – a retrospective study"

### Supplementary Figure 1

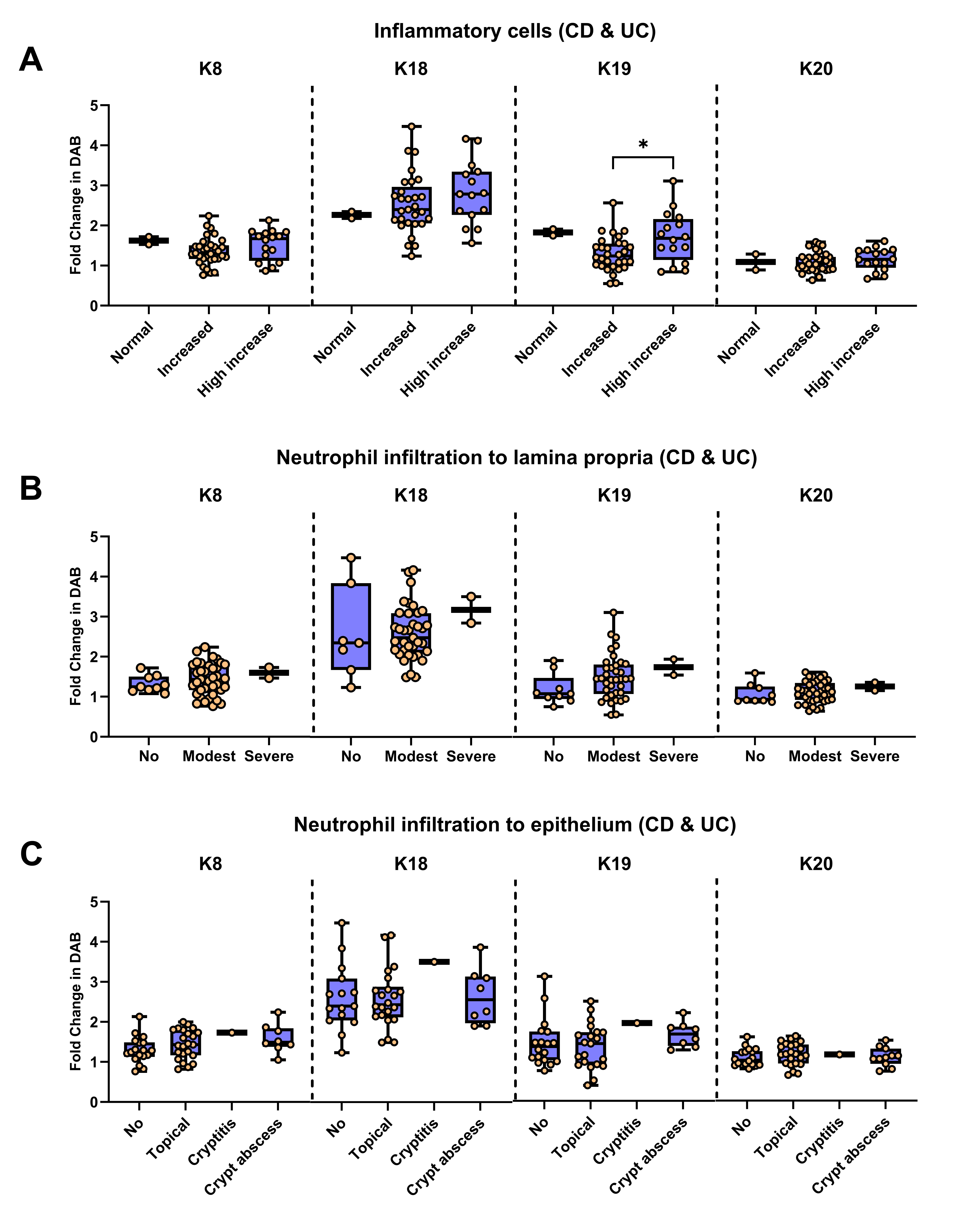
